# Oral delivery of SARS-CoV-2 DNA vaccines using attenuated *Salmonella typhimurium* as a carrier in rat

**DOI:** 10.1101/2020.07.23.217174

**Authors:** Dan Zhu, Yue Meng, Aaodeng Qimuge, Bilige Bilige, Tegexi Baiyin, Temuqile Temuqile, Shana Chen, Siqin Bao, Huricha Baigude, Dezhi Yang

## Abstract

The 2019 novel coronavirus disease (COVID-19) is the disease that has been identified as severe acute respiratory syndrome coronavirus 2 (SARS-CoV-2), but the prophylactic treatment of SARS-CoV-2 is still under investigation. The effective delivery of eukaryotic expression plasmids to the immune system’s inductive cells constitutes an essential requirement for the generation of effective DNA vaccines. Here, we have explored the use of *Salmonella* typhimurium as vehicles to deliver expression plasmids orally. Attenuated *Salmonella phoP* harboring eukaryotic expression plasmids that encoded spike protein of SARS-CoV-2 was administered orally to Wistar rats. Rats were immunized orally with *Salmonella* that carried a eukaryotic expression plasmid once a week for three consecutive weeks. The efficiency of the vaccination procedure was due to the transfer of the expression plasmid from the bacterial carrier to the mammalian host. Evidence for such an event could be obtained in vivo and in vitro. Our results showed that all immunized animals generated humoral immunity against the SARS-CoV-2 spike protein, indicating that a *Salmonella*-based vaccine carrying the Spike gene can elicit SARS-CoV-2-specific humoral immune responses in rats, and may be useful for the development of a protective vaccine against SARS-CoV-2 infection.

## Introduction

Novel coronavirus disease-2019 (COVID-19) is a life-threatening contagious disease caused by severe acute respiratory syndrome coronavirus 2 (SARS-CoV-2). As of 23 June 2020, data from the World Health Organization (WHO) has shown that since the first SARS-CoV-2 case emerged in China’s Wuhan city, in November 2019, it has affected over 8,546,919 cases in over 200 countries and devitalized near 456,726lives. There is no specific and compelling medicine or vaccine that has developed against SARS-CoV-2, currently treated with heteropathy or conservative therapeutics (El-Aziz and Stockand, 2020). Although the first COVID-19 outbreak was controlled by quarantine in China, more and more case reports in other countries indicated that COVID-19 remains a constant threat. Therefore, it is essential to develop a safe and effective vaccine against SARS-CoV-2. Efforts to create safe and effective SARS-CoV-2 vaccines are underway worldwide (Prompetchara et al., 2020). Clinical trials utilizing virus (inactivated or attenuated), viral vector (replicating or non-replicating), nucleic acid (DNA or RNA), and protein-based (protein subunit or virus-like particles) developed across the world (Callaway, 2020; Corey et al., 2020). An inactivated SARS-CoV-2 vaccine seems to be the most facile and convenient option, but inactivated vaccines are more expensive than oral vaccines, considering the scale of the vulnerable population worldwide. Therefore, developing a gene-based vaccine may be a more promising choice (Liu et al., 2005).

The SARS-CoV-2 genome (30 kb in size) encodes a large, non-structural polyprotein (ORF1a/b) that is further proteolytically cleaved to generate 15/16 proteins, four structural proteins and five accessory proteins (ORF3a, ORF6, ORF7, ORF8, and ORF9) (Chan et al., 2020; Li et al., 2020; Wu et al., 2020). Those four structural proteins consist of the spike surface glycoprotein, the membrane protein, the envelope protein, and the nucleocapsid protein, which is essential for SARS-CoV-2 assembly and infection. The spike surface glycoprotein plays a crucial role in its attachment to host cells and can be further cleaved by host proteases into an N-terminal S1 subunit and a membrane-bound C-terminal S2 region (Yuan et al., 2017). Binding of the S1 subunit to a host receptor (i.e., ACE2 receptor) can destabilize the prefusion trimer, leading to shedding of the S1 subunit and transition of the S2 subunit into a highly stable post-fusion conformation (Li et al., 2020). To engage a host receptor, the receptor-binding domain (RBD) of the S1 subunit undergoes hinge-like conformational movements, which transiently hide or expose the determinants of receptor binding (Li et al., 2020; Wrapp et al., 2020).

DNA vaccination, i.e., immunization with an isolated eukaryotic expression plasmid that encodes the antigen rather than immunization with the antigen itself, has added a new dimension to vaccine research (Klinman et al., 2000). The tremendous flexibility offered by DNA vaccination includes the possibility of co-expressing immunomodulatory molecules like cytokines, co-stimulatory molecules, or antisense RNA to steer the immune response into the required direction. Besides, antigens can be manipulated by the addition of sequences that target transport to particular cellular compartments or influence their processing, thus improving the efficacy of immunization. Furthermore, DNA vaccines can be applied in ways that favor either systemic or mucosal responses. Therefore, it is foreseeable that appropriate expression plasmids for a wide variety of applications will be developed to allow a rapid testing of potential protective vaccine candidates in suitable animal models and a quick assessment of optimal strategy (Darji et al., 2000).

A significant drawback of genetic vaccination is its inefficiency, which is inherent in the way that vaccines are currently delivered, and its requirement of relatively large amounts of purified plasmids (Lai and Bennett, 1998). Thus, it is mandatory to develop a carrier system that improves efficacy and possibly target the vaccines *in vivo* inductive sites and cells of the immune system. The use of bacteria as vehicles for genetic vaccination is an attractive and straightforward idea derived from many intrinsic properties afforded by these systems. Previous studies reported that viral expression of the full-length S protein or S1 subunit of SARS coronavirus elicits a high titer of neutralizing antibodies in monkey and rat models (Bukreyev et al., 2004; Gao et al., 2003; Liu et al., 2005). In this study, we used a eukaryotic expression encoding S protein of SARS-CoV-2 by attenuated *Salmonella* carrier and investigated its ability to induce specific antibodies against SARS-CoV-2. Our study demonstrated that specific antibodies against SARS-CoV-2 were elicited in a rat model following the oral administration of SARS-CoV-2-S. This result indicates that our construct could be developed into a safe SARS-CoV-2 vaccine.

## Materials and Methods

### Bacterial strains, growth conditions, and oligonucleotides

*Salmonella enterica* serovar Typhimurium (*S. typhimurium*) strains were derived from the wild-type strain ATCC 14028s (14028). *Salmonella* was grown at 37°C in Luria-Bertani (LB) broth (Difco). When necessary, antibiotics were added to bacterial cultures at final concentrations of 50 μg/ml for ampicillin (Ap), 20 μg/ml for chloramphenicol (Cm), or 50 μg/ml for kanamycin (Km). *E. coli* DH5α was used as hosts for the preparation of plasmid DNA, respectively. Oligonucleotides used in this study are given in Table 1. *Salmonella* phoP (STM1231) was incubated overnight at 37°C with shaking at maximum 50 rpm until an OD600 of 0.6 was reached, then washed and resuspended in phosphate-buffered saline (PBS) with 30% glycerol for oral administration.

**Table 1.**
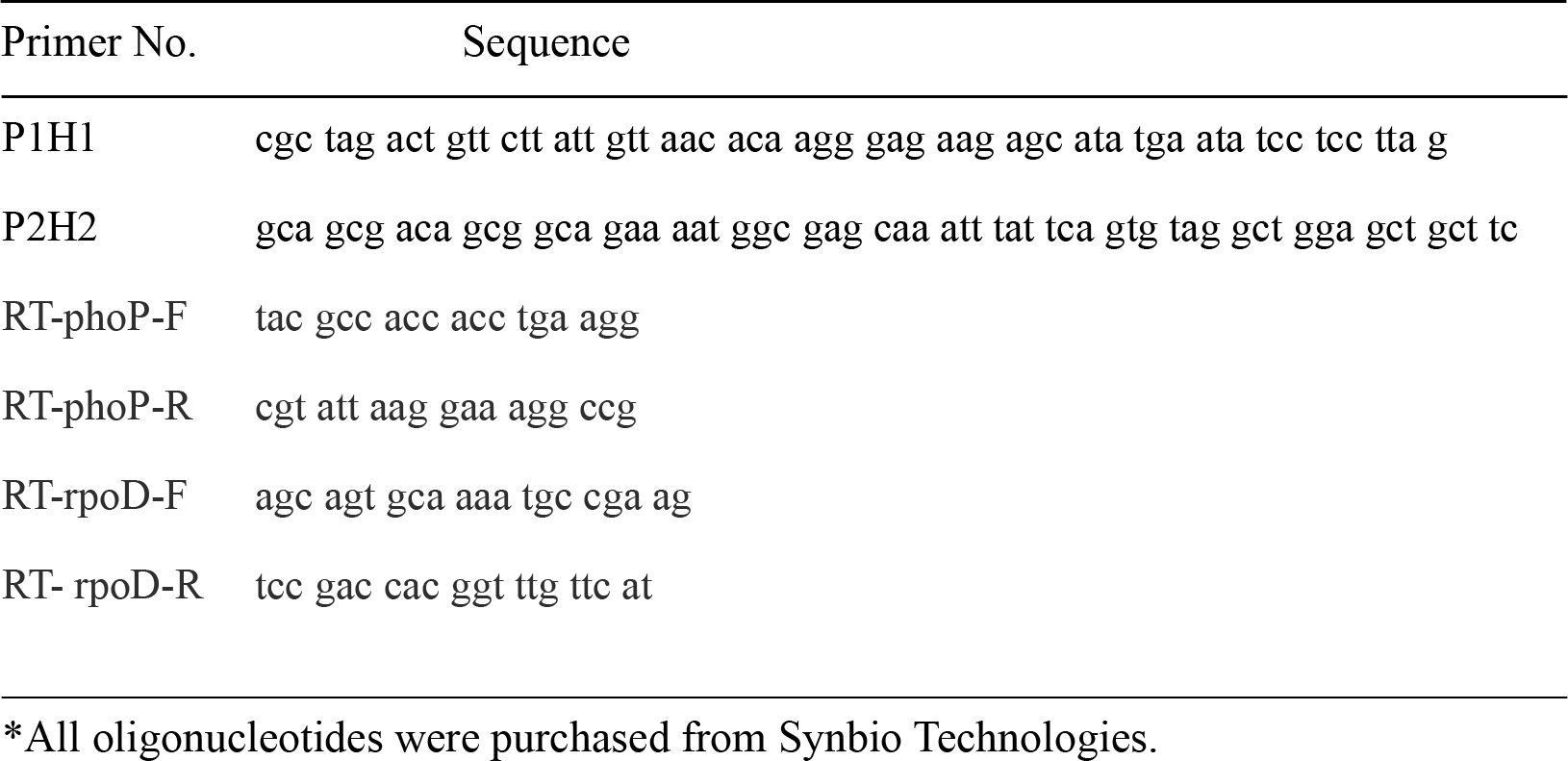
Primers used in this study*.

| Primer No. | Sequence |
| --- | --- |
| P1H1 | cgc tag act gtt ctt att gtt aac aca agg gag aag agc ata tga ata tcc tcc tta g |
| P2H2 | gca gcg aca gcg gca gaa aat ggc gag caa att tat tca gtg tag gct gga gct gct tc |
| RT-phoP-F | tac gcc acc acc tga agg |
| RT-phoP-R | cgt att aag gaa agg ccg |
| RT-rpoD-F | agc agt gca aaa tgc cga ag |
| RT-rpoD-R | tcc gac cac ggt ttg ttc at |

\*All oligonucleotides were purchased from Synbio Technologies.

### Strain construction and transformation of Plasmid into *S. typhimurium* phoP

*Salmonella* strain gene deletion was generated as described previously (Datsenko and Wanner, 2000). Deletions of the *phoP* gene were carried out using primer pairs P1H1 and P2H2, to amplify the kanamycin resistance cassette (KmR) from plasmid pKD4 (Datsenko and Wanner, 2000), the PCR products were electroporated into wild-type cells harboring pKD46, and KmR colonies were selected. The integration of the drug-resistant cassette into the chromosome in these mutants was confirmed by colony PCR. The KmR cassette was removed by using pCP20. Electroporation was carried out by mixing 100 μL of resuspended cells with 0.2 μg plasmid DNA. The suspension was transferred into a disposable cuvette (Bio-Rad Laboratory, Richmond, CA) with an 0.2 cm electrode gap and subjected to an electric pulse at 8 ms, 1.5 kv, 600 Ω and 10 μF using a MicroPulser (Bio-Rad Laboratory, Richmond, CA).

### *In vitro* transfection of pSARS-CoV-2-S plasmid

The pcDNA3.1(+)-CMV-SARS-CoV-2-S-GFP (pSARS-CoV-2-S) plasmid was purchased from Genscript Co., Ltd. (Nanjing, China) (Fig. 1). Plasmid and DNA isolation/purification kits were purchased from TIANGEN Biotech Co., Ltd. (Beijing, China). Transfection reagent DoGOX was described in our previvors study (Ganbold et al., 2017). 293T cells were propagated in Dulbecco’s modified Eagle medium (DMEM) (HyClone, USA) supplemented with 10% fetal bovine serum (Gibco, USA) and maintained in a humidified incubator at 37°C with 5% CO_2_. For the in vitro transfection experiment, pcDNA3.1(+)-GFP control and pcDNA3.1(+)-CMV-2019-nCoV-S-GFP were used. Cells (1.5 × 10^5^ cells/well) were seeded into 6-well plates the day before transfection. Cells were transfected with 2 μg/well of DNA complexed to DoGox for 24 h, as previously described (Xiao et al., 2016).

**Fig. 1.**
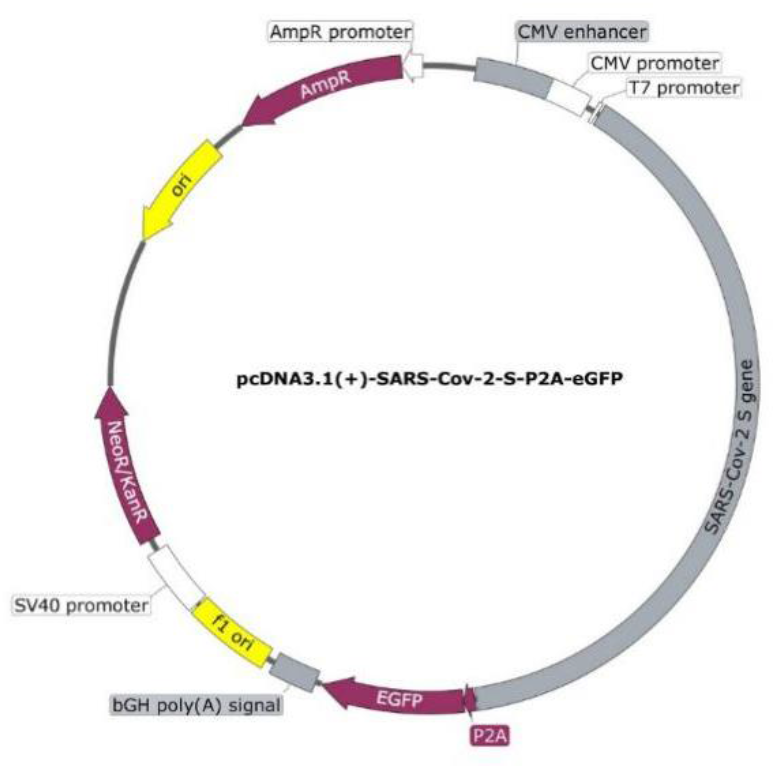
pcDNA3.1(+)-CMV-SARS-CoV-2-S-GFP plasmid map.

**Fig. 2.**
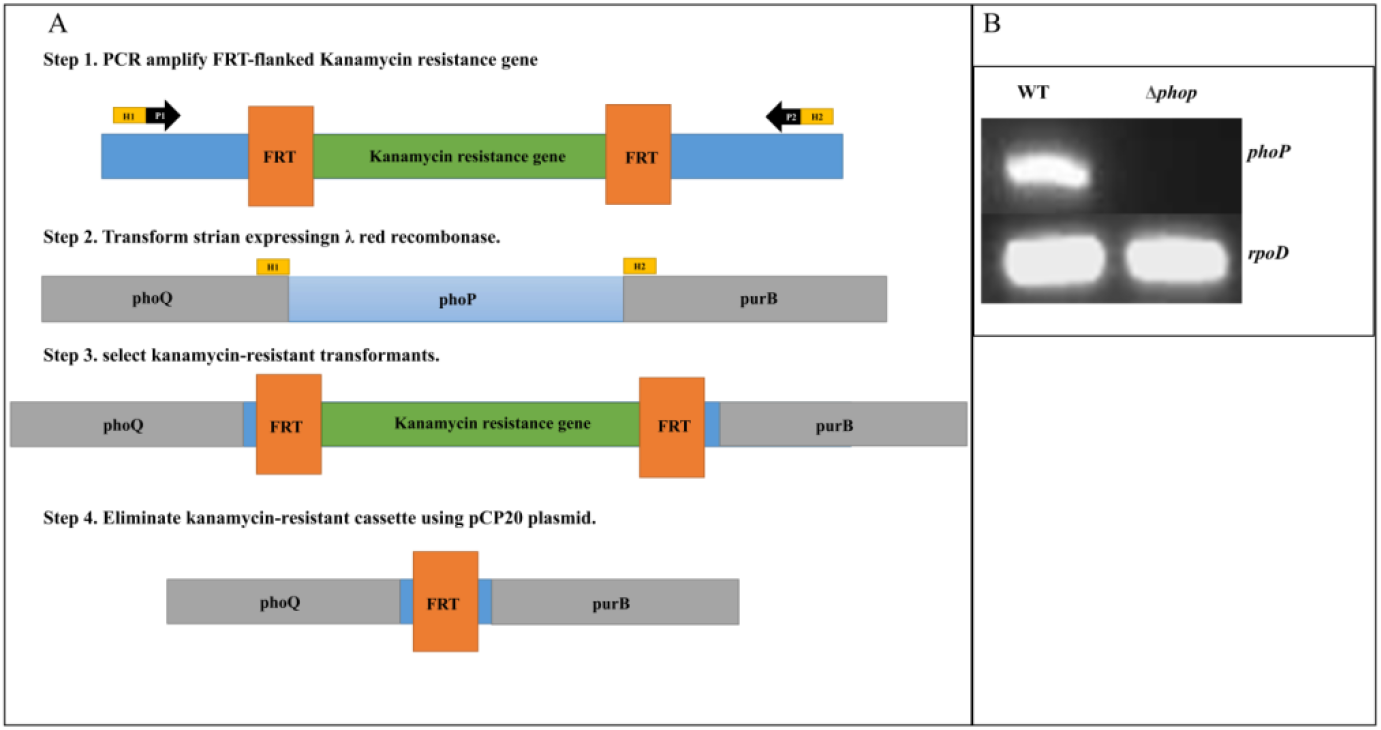
Construction of the *phoP* gene deletion in *Salmonella*. (A) Replacement of the *phoP* gene by kanamycin cassette and removal of the kanamycin cassette by pCP20 plasmid. (B) Genomic DNA PCR test *phoP* gene deletion in wild-type *Salmonella* strain and Δ*phoP* strain.

### Detection of SARS-CoV-2-S protein expression in *vitro*

For the western blot detection of SARS-CoV-2-S protein, the cells were washed twice with ice-cold PBS and harvested from the dishes with a cell scraper by adding a WIP lysis buffer. The recovered lysate was incubated for 30 min on ice and centrifuged at 14,000 x g for 20 min to remove cell debris. The protein concentration of whole-cell lysates was determined by using the BCA Protein Assay kit (Pierce). Protein samples were separated by 8% SDS-PAGE and transferred to PVDF membranes (Amersham), then proteins were monitored with ECL Western blotting substrate (Pierce) after incubation with anti-SARS-CoV-2-S-RBD (Sino Biological, China) or anti-β-actin antibody (Solarbio, China).

### *SARS-CoV-2-S* DNA immunization delivered by attenuated *S. typhimurium*

A total of 18 female Wistar rats were randomly divided into two groups, with nine rats each. The first group was immunized orally with 1×10 ^7^ attenuated *S. typhimurium* phoP transformed with pcDNA3.1-GFP in 200 μl of PBS. The second group of rats was given 200 μL of PBS with 1×10 ^7^ attenuated *S. typhimurium* PhoP transformed with pcDNA3.1(+)-CMV-SARS-CoV-2-S-GFP. All mice were given 200 μL of 10% NaHCO_3_ orally to neutralize gastric acids before oral inoculation with bacteria. All groups of the rat were boosted three times with the same vaccine components at two weeks interval.

### Enzyme linked immunosorbant assay (ELISA)

As previously described, the sera were used to test the anti-SARS-CoV-2-S IgG antibodies by enzyme-linked immunosorbent assay (ELISA). (Vedi et al., 2008). Briefly, ELISA was performed in 96-well plates (Costar, USA) coated with 0.5 μg of rSARS-CoV-2-S (KITGEN BIOTECHNOLOGY, China) per well in 100μl bicarbonate buffer (pH 9.6). After being blocked with 5% FCS, 100 μl serum samples at several dilutions in PBS were added to the wells and incubated at 37°C for one hour. Then the HRP conjugated rabbit anti-goat IgG antibodies (1:3000, Solarbio, China) were added, and incubation continued at 37°C for one hour. The plate was developed using TMB (Solarbio, China), following 2 M H_2_SO_4_ addition to stopping the reaction, and read at 450 nm by ELISA plate reader for final data. Samples were analyzed in duplicate.

### Statistical analyses

Statistical analyses were performed with one-way ANOVA using GraphPad Prism 6.0. All data were expressed as the mean ± standard deviation.

## Results

### Construction of *S. Typhimurium* strains with *phoP* mutation

Given the ultimate objective of constructing a vaccine strain that would be safe in mammals, we generated a Δ*phoP* strain by λ-Red recombination engineering using the one-step inactivation procedure developed by Datsenko and Wanner using pKD4-amplified PCR products. λ-Red functions were expressed from plasmid pKD46. Kanamycin cassettes were excised with the pCP20 plasmid as previously described (Fig. 1A-B) (Datsenko and Wanner, 2000). Strains were verified by genomic DNA PCR by using primers pair for the *phoP* gene with RT-phoP-F& RT-phoP-R or *rpoD* gene with RT-rpoD-F& RT-rpoD-R. PCR result indicate the *phoP* gene was removed from the chromosome site.

### Expression of the recombinant eukaryotic expression plasmid pcDNA3.1(+)-CMV-SARS-CoV-2-S-GFP in 293T cells

In order to investigate the ability of pcDNA3.1(+)-CMV-SARS-CoV-2-S-GFP to express the SARS-CoV-2-S protein, transfection, and western blot analysis were performed. A P2A ribosomal self-skipping sequence separates SARS-CoV-2-S from a GFP marker. The strong CMV promoter was driving SARS-CoV-2-S-GFP co-transcription and the P2A cleavage site separation of SARS-CoV-2-S and GFP during translation in eukaryotic cells (Daniels et al., 2014). The results demonstrated that 293T cells transfected with pcDNA3.1(+)-CMV-SARS-CoV-2-S-GFP expressed the SARS-CoV-2-S protein compared wit the cells transfected with the empty pcDNA3.1(+)-CMV-GFP plasmid (Fig. 3A–B).

**Fig. 3.**
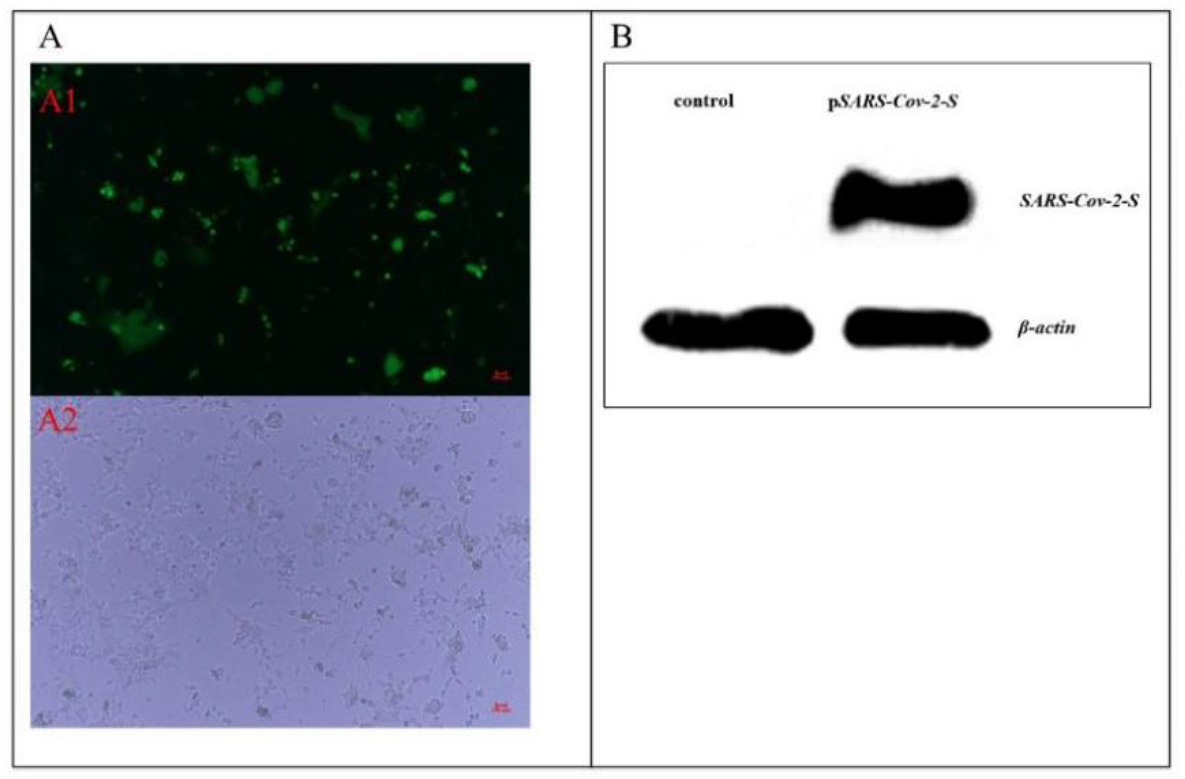
Micrographs of 293T cells transfected with pSARS-CoV-2-S (X 100). (A1, A2) 293T cells transfected with pSARS-CoV-2-S (pcDNA3.1(+)-CMV-SARS-CoV-2-S-GFP) at 48 hours after transfection. A1 fluorescence micrograph with GFP expression in cells and light micrograph with the same visual field as A2. (B) 293T cells transfected with pSARS-CoV-2-S at 48 hours after transfection. The SARS-CoV-2-S protein showed about 141 kDa.

### Anti-SARS-CoV-2-S protein sera IgG responses in the rat after orally *Salmonella*-mediated DNA vaccination

Rats (n = 9 per group) were orally immunized with Δ*phoP* strain with pSARS-CoV-2-S (1 × 10^7^ cfu/(dose·rat)), or Δ*phoP* strain with controls (plasmid) (Fig. 4). Rats were immunized on days 0, 7, and 14. Sera were collected on days at 0, 14, and 28 days post-immunization (d.p.i). Estimate the antigen-specific humoral immune response after immunization with pSARS-CoV-2-S, antibody ELISA was performed on the sera collected from the rat immunized with pSARS-CoV-2-S or control plasmid. The pSARS-CoV-2-S vaccine-elicited humoral immune response, as evidenced by the fold increased (pSARS-CoV-2-S group at 14 days, three rats of the nine rats increased to 4-5 times and another six rats increase to 8-9 times when compared with the control group. At 28 days, all rats increase to 4-6 times when compared with the control group, see Fig. 5) in the OD values between pSARS-CoV-2-S and control groups (None increase in all experiment period). This profile of IgG suggested that the novel consensus oral vaccine strategy used in this study elicited an antigen-specific humoral immune response (Fig. 5).

**Fig. 4.**
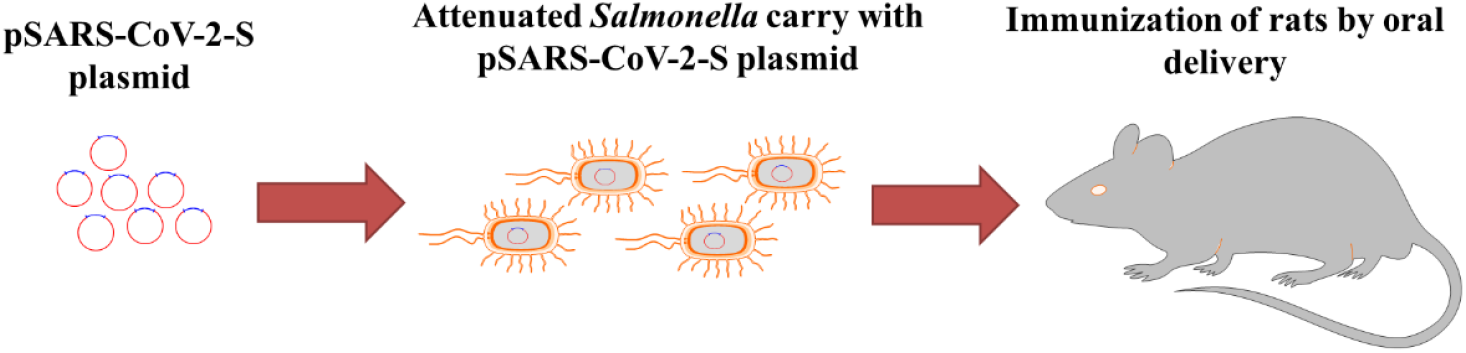
The procedure of plasmid transformation to *Salmonella* strain and immunization of rat.

**Fig. 5.**
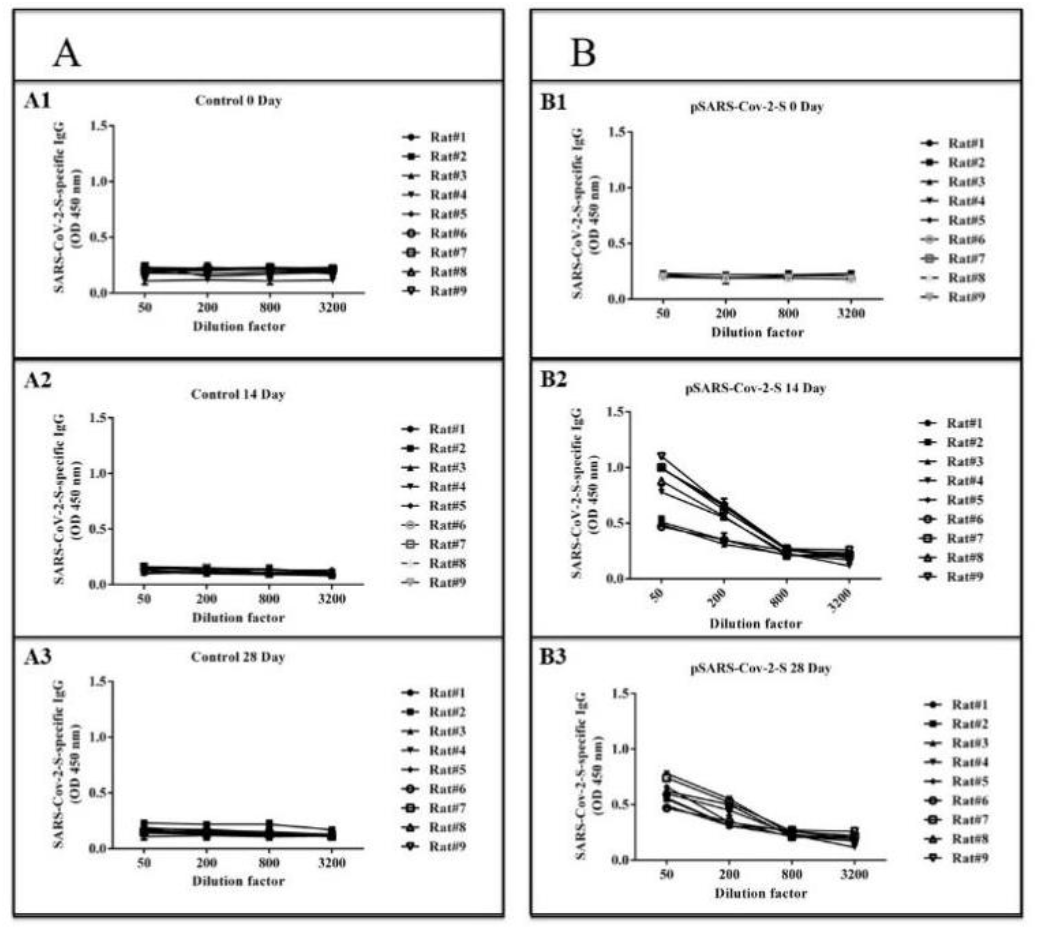
Humoral responses to SARS-CoV-2-S protein antigen in the rat after immunizationon day 0, day 14, and day 28 with *Salmonella* carrying the control vector or pSARS-CoV-2-S (as described in the methods). (A) After immunization with the control vector, test SARS-CoV-2-S protein antigen binding of IgG in serial serum dilutions from a rat at day (0, 14, 28). Data shown represent test mean OD450 nm values (mean±SD) for each of 9 rats, or (B) After immunization with the pSARS-CoV-2-S vector, SARS-CoV-2-S protein antigen binding of IgG in serial serum dilutions from a rat at day (0, 14, 28). Data shown represent mean OD450 nm values (3 times measurement, mean±SD) for each of 9 rats.

## Discussion

Various strategies have been explored to search for an effective vaccine against SARS-CoV-2, including the use of inactivated whole virus particles, attenuated live virus and gene vaccines (Gao et al., 2020; Smith et al., 2020; van Doremalen et al., 2020; Wang et al., 2020; Zhu et al., 2020). Recently, the the gene vaccines that express SARS-CoV structural proteins and induce host immunity against SARS-CoV-2 have become increasingly popular.

Several bacterial species have been used to transfer eukaryotic expression plasmids into host cells. *Shigella flexneri*, *Listeria monocytogenes*, and recombinant *Escherichia coli* that harbor the invasion plasmid of *Shigella* have been used successfully (Gentschev et al., 2000; Schoen et al., 2004; Sizemore et al., 1995). Since these bacteria escape from the phagosome, it has been postulated that entry into the cytosol would be a crucial step in bacteria mediated transfection. Using attenuated strains of *S. flexneri* and *S.typhimurium* as DNA carrier, antibody responses against the transgene could be detected in the animal model (Darji et al., 2000; Fennelly et al., 1999).

Although oral administration of the vaccine is challenging, the socio-economic benefits of such vaccination are apparent. Moreover, oral vaccination could stimulate both humoral and cellular immune responses (De Smet et al., 2014). The most successful oral vaccine are the polio vaccine (Smith and Leggat, 2005). Our design of pSARS-CoV-2-S plasmid-based orally delivered vaccine was able to induce significant antigen-specific IgG antibodies. As indicated earlier, the COVID-19 protein SARS-CoV-2-S is considered a robust putative vaccine target due to it being significantly associated with the stimulation of SARS-CoV-2 (Smith et al., 2020). This protein was found to be immunogenic and capable of eliciting a humoral response. Since the previous study developed a vaccine against MERS coronavirus; and other published studies of SARS vaccines, S protein was chosen as the antigen target (Modjarrad et al., 2019; Smith et al., 2020; Wang et al., 2020). The SARS-CoV-2 S protein is a class I membrane fusion protein, which the major envelope protein on the surface of coronaviruses. Initial studies have already been performed, which indicate antibodies can block SARS-CoV-2 interaction with its host receptor (ACE2) (Zhou et al., 2020). In vivo immunogenicity studies show the functional antibodies and T cell responses in multiple animal models after SARS-CoV-2-S expression plasmid or mRNA direct immunized to the animals or human (Smith et al., 2020; Wang et al., 2020). The oral administration of the SARS-CoV-2-S oral vaccine was found to induce the response. In the following study, the protection efficiency will be evaluated with a high dose of viral challenge in nonprimate mammals such as rhesus monkey. In conclusion, attenuated *Salmonella typhimurium* carrying plasmid encoding SARS-CoV-2-S is one of the promising candidates for developing an oral delivery vaccine against SARS-CoV-2 infection in humans.

## Acknowledgments

This research was financially supported by screening of the Mongolian medicine Treatment of COVID-19 in 2020, Inner Mongolia, China (T101499.404). The authors are grateful to Guojun Wang and Digengni for language revision. Dr. Morigen is acknowledged for share plasmid pKD4, pKD46, pCP20. Dr. Haisheng Yu is acknowledged for share plasmid pcDNA3.1(+)-CMV-SARS-CoV-2-S-GFP.

## Conflict of interest

No conflict interest declared.

## Author Contributions

1. **Conceived and designed the experiments:** DY BB TT.
2. **Performed the experiments:** DZ YM AQ.
3. **Analyzed the data:** DZ YM DY.
4. **Contributed reagents/materials/analysis tools:** TB BB.
5. **Wrote the paper:** HB DY.

## Notes

### Competing Interest Statement

The authors have declared no competing interest.

